# Three’s a crowd: The relationship among endoparasites, an epibiont and their *Daphnia* host

**DOI:** 10.1101/2024.06.03.597088

**Authors:** Ofir Hirshberg, Frida Ben-Ami

## Abstract

In freshwater communities, organisms interact in a variety of ways, including predation, competition and parasitism. Parasites are ubiquitous, playing an important role in shaping freshwater communities. Endoparasites live within internal organs of their host, while ectoparasites, also known as epibionts, are confined to the external part of the host’s body. We conducted a series of experiments to examine the relationship between endoparasites and epibionts using the crustacean *Daphnia magna* as host, the rotifer *Brachionus rubens* as epibiont and three species of endoparasites. First, we tested host preference of the epibiont between *Daphnia* infected by endoparasites and uninfected *Daphnia*. Epibiont were found to attach more to uninfected *Daphnia* than to *Daphnia* infected by the yeast *Metschnikowia bicuspidata.* On the other hand, epibionts attached more to *Daphnia* infected by the microsporidium *Hamiltosporidium tvaerminnensis* than to uninfected *Daphnia*. Second, we examined the effect of epibionts on the infection of *Daphnia* by endoparasites. Infection prevalence tended to be higher, though not significantly, in the presence of epibionts. For two of the endoparasites, *M. bicuspidata* and *H. tvaerminnensis*, infection intensity (i.e., parasite spore production) was higher in the presence of epibionts. The infection intensity of *M. bicuspidata* and the bacterium *Pasteuria ramosa* was affected by the time of death of the *Daphnia* (i.e., virulence). Finally, we examined the effects of endoparasites and epibionts on the survival and offspring production of the *Daphnia*. Both host survival and offspring production were negatively affected by the endoparasites, while epibionts did not seem to affect the fitness of their host.

## Introduction

In natural communities, organisms interact in a variety of ways, including predation, competition and parasitism. These biotic interactions drive ecological and evolutionary processes and thus play a key role in the creation and maintenance of biodiversity and ecosystem functioning. Biotic interactions between two organisms are diverse and may vary among different groups of organisms. They can be beneficial to both organisms (e.g., mutualism), but are usually harmful to at least one of them, as in the case of parasitism, which is extremely common in natural communities. Several estimates suggest that at least 50% of plants and animals are parasitic at some stage during their life cycle (Bush, 2001). Many studies have shown that parasites play a key role in shaping natural communities, as they reduce the survival and fecundity of their host (Decaestecker et al., 2005; Stirnadel and Ebert, 1997), thereby affecting its population size. Additionally, parasites may also indirectly affect host population by altering mate choice and predation risk (Goren and Ben-Ami, 2017; Thomas et al., 1997).

Parasites are highly diverse in their taxonomic affiliation (Weinstein and Kuris, 2016). While some parasites are specialists, infecting only a small number of host species or genotypes, others are generalists, infecting a large number of hosts (Stirnadel and Ebert, 1997). The ability of parasites to infect their host largely depends on physiological and environmental characteristics (Lafferty and Kuris, 1999). Furthermore, parasitic infections rarely occur in isolation, with many infections involving two or more parasite strains (Balmer and Tanner, 2011) or species (Bordes and Morand, 2011; Halle et al., 2024). Interactions between these parasites can be positive when infection by one parasite enhances the transmission or infection by another, or negative when one parasite suppresses another (Dallas et al., 2019).

Endoparasites reside within their host’s body. They live and reproduce in different parts of their host’s body (Dallas et al., 2019) and have different physiological and behavioral influences (Thomas et al., 1997). Endoparasites include all viruses, many bacteria and some eukaryotes. They are common in freshwater zooplankton communities, belong to diverse taxonomic groups and infect a variety of hosts (Goren and Ben-Ami, 2013), and are capable of reducing their host population size (Ebert et al., 2000).

Ectoparasites, also known as epibionts, are confined to the exterior part of their host’s body. While they were found to be less damaging to their host than endoparasites (Decaestecker et al., 2005), fitness reduction and a decrease in fecundity and survival as a result of epibionts have been reported (Green, 1974; Pauwels et al., 2014). Other epibionts were found to induce injuries and hair or feather loss (Lehmann, 1993). The epibiont community living upon freshwater crustaceans is highly dependent on host molting, as molting crustaceans shed their epibionts, creating a clean substrate for new epibionts. Some epibionts of freshwater zooplankton are generalists, being found on a wide range of substrate organisms, while others are specialist, confined to a single species or genus (Threlkeld et al., 1993). Epibionts can be found both solely on substrate organisms as well as swimming freely in the water column or attached to a non-moving substrate.

The present study investigates the relationship between endoparasites and epibionts, using the water flea *Daphnia magna* (Cladocera: Daphniidae) as the host, the rotifer *Brachionus rubens* (Ploima: Brachionidae) as the epibiont and three endoparasites belonging to different taxonomic groups. First, we conducted several short-term experiments designed to examine host preference of the epibiont between uninfected hosts and hosts infected by each of the endoparasites. Given that *B. rubens* was shown to attach in different quantities to different host species as well to males vs. females (Hirshberg and Ben-Ami, submitted), we hypothesized that the epibiont would attach in different quantities to infected and uninfected hosts. Second, we examined the effects of the epibiont on *D. magna* infected by each of the endoparasites, considering both infection prevalence and infection intensity (i.e., parasite spores produced). Since endoparasites rely on their host’s resources just like epibionts, rotifers (our epibiont) are expected to compete with their host over food (Pauwels et al., 2014). Thus, we hypothesized that infection by the epibiont would suppress infection by endoparasites, by decreasing the availability of host resources. Finally, we examined the combined effects of the epibiont and endoparasites on the survival and offspring production of their host. We hypothesized that such combination would increase parasite-induced host mortality.

## Methods

### Study organisms and cultures

#### Host

The water flea *Daphnia magna* (Branchiopoda: Cladocera) is among the largest and most common freshwater crustacean. Populations of *D. magna* are usually composed of parthenogenetic females that reproduce asexually, producing clonal offspring. However, as a result of various environmental cues, *D. magna* produce males and reproduce sexually (Hobaek and Larsson, 1990). *D. magna* is host to a variety of parasite species, including viruses, fungi, bacteria and microsporidia (Ebert, 2005). This, along with its quick asexual reproduction lifecycle and small size, make *D. magna* an excellent model organism for studying host-parasite interactions and coevolution. The ILBM1 *D. magna* clone used in our study originated from a small ephemeral pond in the Negev desert in Israel. Clonal culture of *D. magna* was established in a climate room at 20°C and a light regime of 12:12 L:D, and fed every other day with 200×10^6^ cells of the algae *Scenedesmus gracilis*.

#### Epibiont

The monogonont rotifer *Brachionus rubens* (Ploima: Brachionidae) is a widespread epibiont that attaches to the body surface of large aquatic invertebrates. *B. rubens* can attach to a non-moving substrate as an epibiont or swim freely in the water column. Increasing the risk of predation was shown to induce attachment of *B. rubens* (Gilbert, 2019). *B. rubens* was also shown to decrease the growth rate and population density of several *Daphnia* species (Iyer and Rao, 1993). Clonal culture of *B. rubens* originated from the Aegelsee pond in Switzerland, was cultivated in a climate room at 20°C and a light regime of 12:12 L:D, and fed every other day with 100×10^6^ cells of the algae *Scenedesmus gracilis*.

#### Endoparasites

The yeast *Metschnikowia bicuspidata* (Ascomycota: Saccharomycetales) is a generalist endoparasite, infecting several species of *Daphnia* (Ebert, 2005). Infection occurs when a mature, needle-shaped spore of *M. bicuspidata* pierces the gut wall of its filter-feeding host before reaching the hemolymph. *M. bicuspidata* usually kills its host within 2-3 weeks post infection. After the host dies, new mature spores are released to the environment. The *M. bicuspidata* strain used in our experiments originated from Lake Vir in Croatia and was propagated using *D. magna* as host.

The bacterium *Pasteuria ramosa* is an obligate parasite of *Daphnia* (Ebert et al., 1996). Infection occurs through the digestion of spores by the filter-feeding host. After infection the bacterium goes through several developmental stages, ending in the formation of millions of mature spores that are released into the waterbody upon host death (Ebert et al., 2016)*. P. ramosa* usually kills its daphniid host within 30-50 days. The clone used in our experiments originated from a pond in Gaarzerfeld, Germany and was propagated using *D. magna* as host. Additionally, some infected *D. magna* were preserved at −20°C before extracting the *P. ramosa* spores.

The microsporidium *Hamiltosporidium tvaerminnensis* is transmitted both horizontally and vertically, as spores are either released to the environment from the decaying body of a dead host, or transferred from mother to offspring. Vertical transmission can be 100% effective for parthenogenetic offspring (Routtu and Ebert, 2015). The G3 isolate used in our experiments was obtained from the Shafir pond, an ephemeral pond in Israel, and propagated using *D. magna* as host.

### Experimental design

#### Experiments 1-3: Host preference

In experiments 1-3, we tested the host preference of the epibiont *B. rubens* between uninfected *D. magna* and *D. magna* singly infected with each of three species of endoparasites. We filled 70 jars with 100 ml of artificial *Daphnia* medium (ADaM, Klüttgen et al., 1994; Ebert et al., 1998). A pair of similar-sized *D. magna* was placed in each jar (*Daphnia* size was measured before they were transferred into the experimental jars). One of the *Daphnia* was uninfected and the other infected by one of three species of endoparasites as per Table 1. Since moving the *Daphnia* using a pipette can cause stress, which may affect their swimming behavior and thus influence the results of the experiment, *D. magna* were acclimated in the experimental jars 24 hours before the beginning of the experiments.

**Table 1:** List of all experiments conducted, including initial number of epibionts and endoparasitic spores (i.e., parasite dose).

| Experiment | Research question | Endoparasite | Number of *Daphnia* (sample size) | Initial number of *B. rubens* | Initial dose of endoparasitic spores | Experiment duration |
| --- | --- | --- | --- | --- | --- | --- |
| 1 | Host preference of *B. rubens* between infected and uninfected *D. magna* | *M. bicuspidata* | 32 pairs | 20 | NA | 6 hours |
| 2 | Host preference of *B. rubens* between infected and uninfected *D. magna* | *P. ramosa* | 46 pairs | 20 | NA | 6 hours |
| 3 | Host preference of *B. rubens* between infected and uninfected *D. magna* | *H. tvaerminnensis* | 53 pairs | 20 | NA | 6 hours |
| 4 | Effects of *B. rubens* on endoparasite infection and fitness of *D. magna* | *M. bicuspidata* | 142 | 40 | 10,000 | 40 days |
| 5 | Effects of *B. rubens* on endoparasite infection and fitness of *D. magna* | *P. ramosa* | 130 | 40 | 100,000 | 40 days |
| 6 | Effects of *B. rubens* on endoparasite infection and fitness of *D. magna* | *H. tvaerminnensis* | 123 | 40 | 350,000 | 40 days |

In the beginning of the experiments, 20 *B. rubens* individuals were introduced to each jar. The jars were then kept in a climate room with a temperature of 20^0^C. After six hours each *Daphnia* was collected, placed in an Eppendorf tube and preserved in Lugol solution. Preliminary experiments have shown that six hours are sufficient for *B. rubens* to attach to the *Daphnia* and that *B. rubens* remains attached after being preserved in Lugol. The *B. rubens* individuals attached to each *Daphnia* were counted under a Leica M205C dissecting microscope. Thereafter, the *Daphnia* were crushed and scanned for endoparasitic spores under a Leica 2500 phase contrast microscope. Jars in which one of the *Daphnia* had died, molted or produced offspring were excluded from the analysis.

#### Experiments 4-6: Epibiont-endoparasites interactions

In experiments 4-6, we tested the effect of the epibiont *B. rubens* on the infection of *D. magna* by each of three species of endoparasites. In addition, we tested the combined effects of the epibiont and endoparasites on host survival and offspring production. To this end, we filled 40 jars with 20 ml of ADaM. A single 1-day-old *Daphnia* was placed in each jar, after which 40 *B. rubens* individuals were inserted into the jar. After 5 days, when the *Daphnia* were covered by the epibionts, spores of each endoparasite were placed in each jar as per Table 1. After 48 hours, *Daphnia* were transferred into new jars with 100 ml of ADaM. During the experiments, *Daphnia* mortality was monitored every day and its offspring were counted every other day. *Daphnia* that had died on the first week of the experiment were removed from the analysis, as their mortality is unlikely due to the effects of the epibiont and/or endoparasites. The remaining dead *Daphnia* were preserved in 20% glycerol for further analysis. After 40 days, all remaining *Daphnia* were preserved in 20% glycerol. Three control groups were established for each experiment. One group was exposed to the endoparasite without the epibiont, the second group was exposed to the epibiont without the endoparasite, and the third group was not exposed to either epibiont or endoparasites. Due to the decline of the *B. rubens* population during the experiment, they were renewed twice, after ten and thirty days. After the experiment each *Daphnia* was crushed and parasite spores were counted using a Thoma counting chamber under a Leica 2500 phase contrast microscope.

### Data analysis

In experiments 1-3, in order to test if the number of rotifers that attached differed between infected and uninfected *D. magna*, we used a-parametric Wilcoxon test for paired data, since the data did not meet the assumption of normality. In experiments 4-6, in order to test if infection prevalence differed among endoparasites and between hosts with and without epibionts, we used binary logistic regression followed by ANOVA. In order to test the effects of epibionts on the infection intensity of endoparasites (i.e., parasite spores produced), we used ANCOVA with time-until-host-death as a covariant. Since the number of spores of *H. tvaerminnensis* did not meet the assumption of equal variances, we applied the log transformation before running the ANCOVA. To further examine the effect of time-until-host-death on infection intensity, we used linear regression. To test the effects of the epibiont and endoparasites on offspring production of *D. magna*, we used two-way ANOVA. Since our data did not meet the assumptions of normality and equal variances, we applied the log(x+1) transformation before running the ANOVA. In order to test for differences in offspring production and time until host death between the test (infected) and control groups, we used pairwise Wilcoxon test. All analyses were performed using R version 4.1.1.

## Results

In experiment 1, the size of infected *Daphnia* ranged between 1.6-4.6mm^2^ and the size of uninfected *Daphnia* ranged between 1.9-4.8mm^2^. The number of epibionts attached to *Daphnia* infected by *M. bicuspidata* ranged between 0-13 with a mean±SE of 4.52±0.54, significantly lower than the number of epibionts attached to uninfected *Daphnia*, which ranged between 1-17 with a mean±SE of 6.26±0.68 (Paired Wilcoxon V=123, p=0.025; Figure 1, left panel). In experiment 2, the size of infected *Daphnia* ranged between 4.5-6.5mm^2^ and the size of uninfected *Daphnia* ranged between 3.9-6.7mm^2^. The number of epibionts attached to *Daphnia* infected by *P. ramosa* ranged between 0-11 with a mean±SE of 3.46±0.47, and did not differ from the number of epibionts attached to uninfected *Daphnia*, which ranged between 0-12 with a mean±SE of 3.24±0.48 (Paired Wilcoxon V=480, p=0.726; Figure 1, middle panel). In experiment 3, the size of infected *Daphnia* ranged between 2.4-6.6mm^2^ and the size of uninfected *Daphnia* ranged between 2.4-6.1mm^2^. The number of epibionts attached to *Daphnia* infected by *H. tvaerminnensis* ranged between 0-16 with a mean±SE of 4.02±0.53, significantly higher than the number of epibionts attached to uninfected *Daphnia*, which ranged between 0-10 with a mean±SE of 2.96±0.38 (Paired Wilcoxon V=749, p=0.049; Figure 1, right panel).

**Figure 1:**
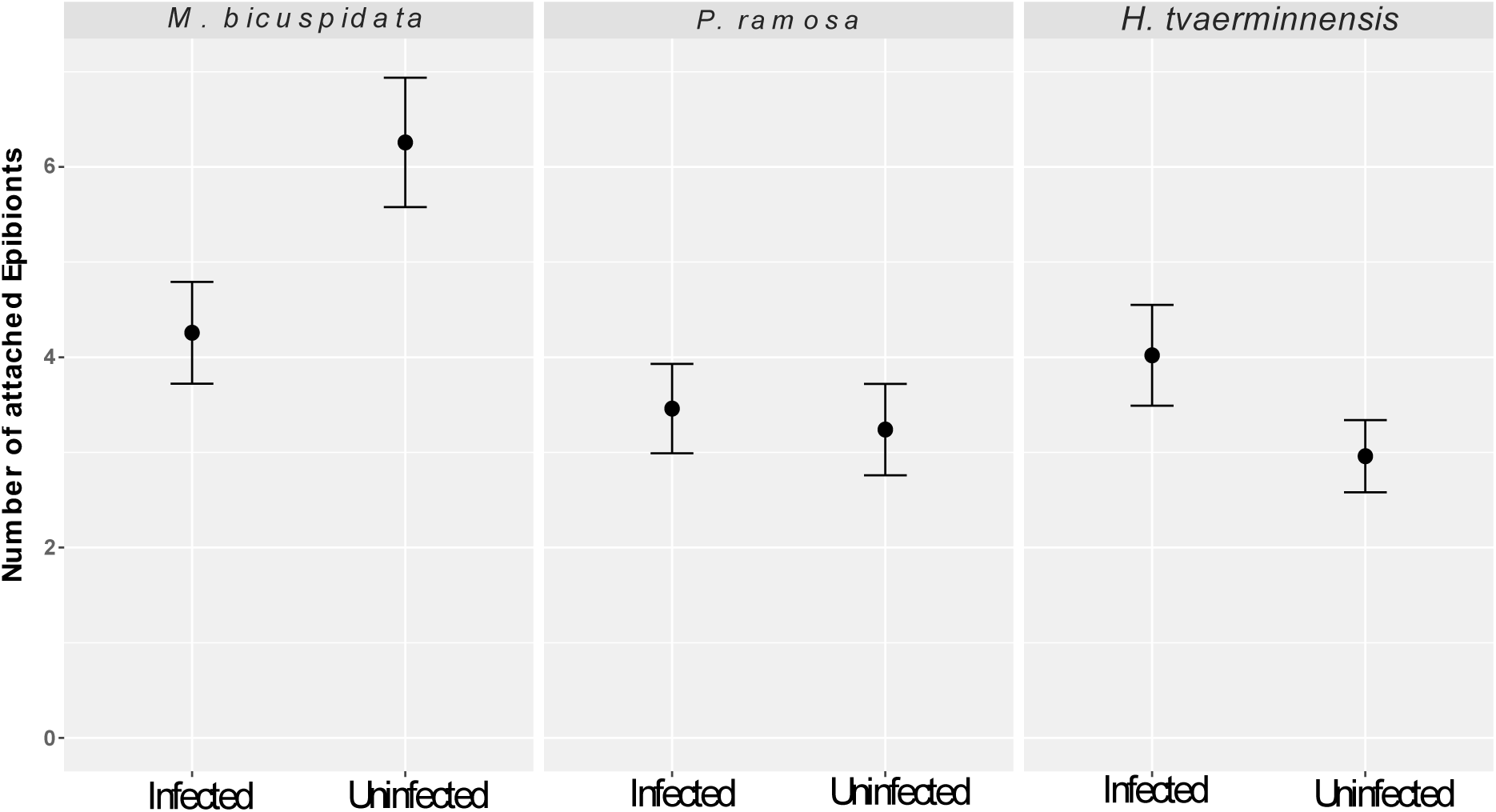
Mean±SE of the number of epibionts attached to *Daphnia* infected by each endoparasite species vs. uninfected *Daphnia*.

In experiments 4-6, the prevalence of infection differed among endoparasites (ANOVA F=19.13, p<0.001; Figure 2). It was highest in *Daphnia* exposed to *M. bicuspidata* and lowest in *Daphnia* exposed to *H. tvaerminnensis.* For all endoparasites, the prevalence of endoparasitic infection was higher in the presence of the epibiont *B. rubens,* though this difference was not significant (ANOVA F=3.5, p=0.063; Figure 2).

**Figure 2:**
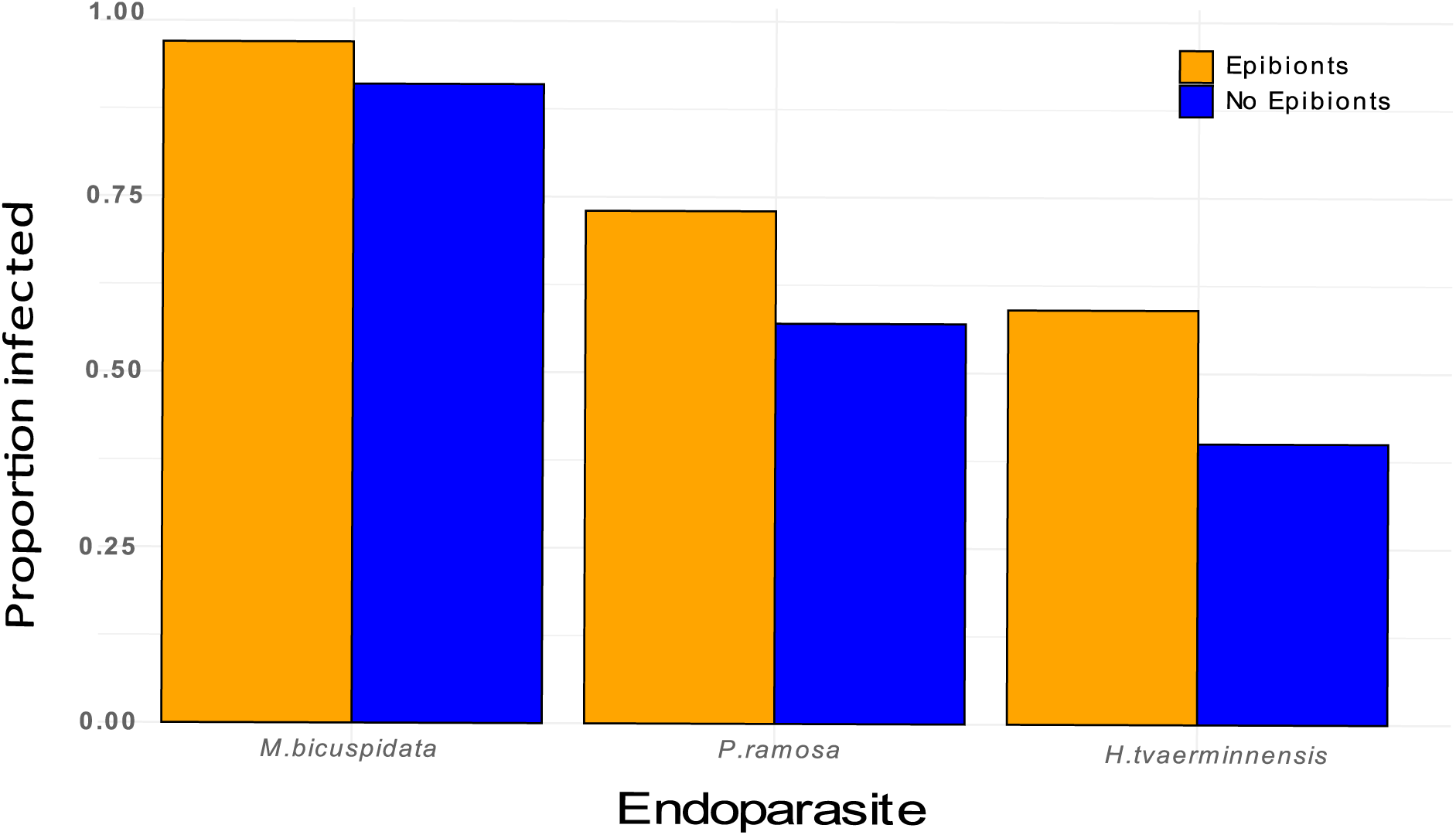
Infection prevalence of three species of endoparasites in the presence and absence of epibionts.

Infection intensity (parasite spores produced) of *M. bicuspidata* was significantly higher in the presence of the epibiont *B. rubens*. Furthermore, the longer it took the *Daphnia* to die, the higher was the intensity of *M. bicuspidata* infection. The intensity of infection by *P. ramosa* was not affected by the presence of epibionts, but it was positively affected by time-until-host-death. Infection intensity of *H. tvaerminnensis* was significantly higher in the presence of epibionts, but it was not affected by time-until-host-death (Table 2; Figure 3).

**Figure 3:**
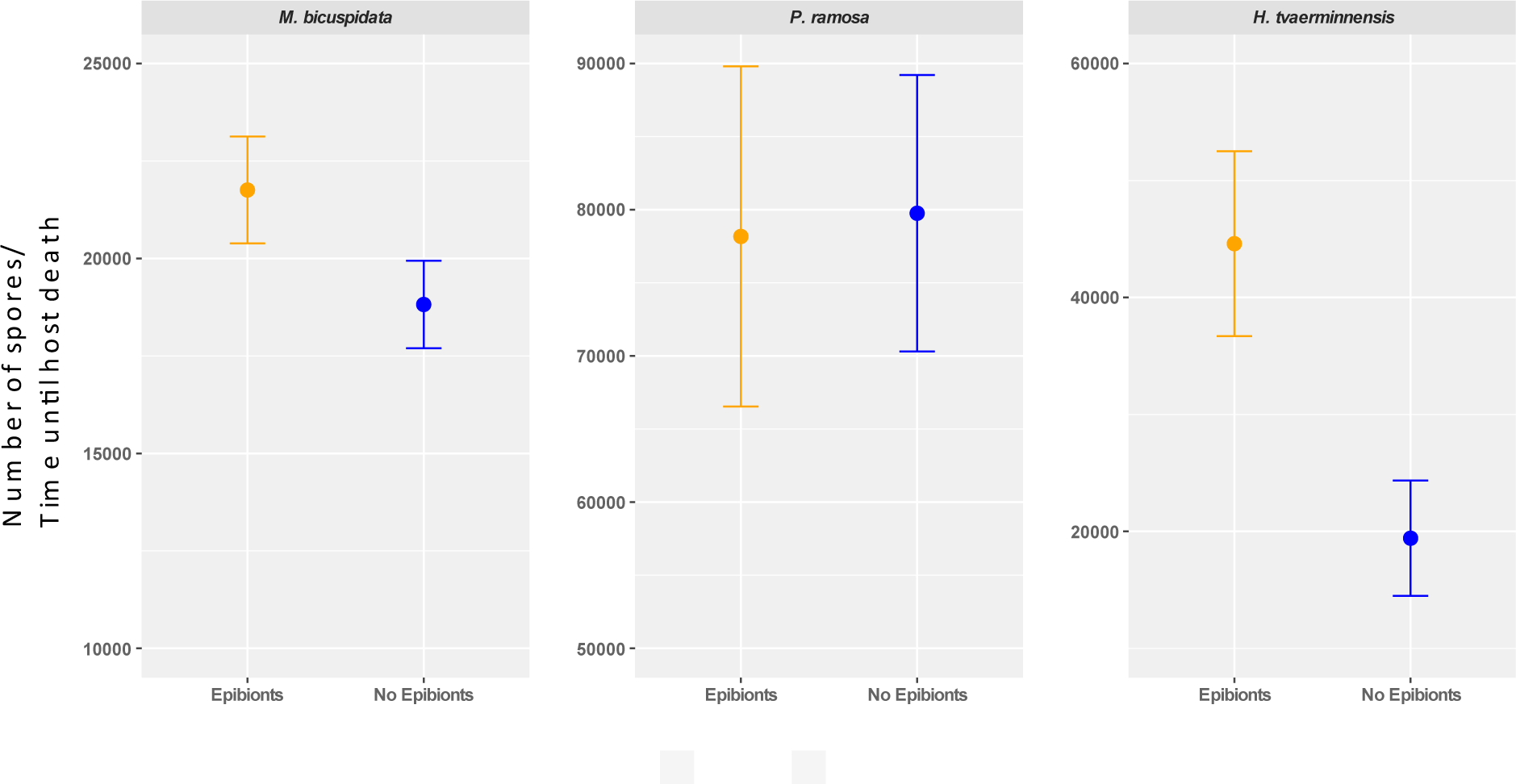
Infection intensity (number of parasite spores), divided by the time-until-host-death, of each of the three endoparasites in the presence and absence of epibionts.

**Table 2:** Results of ANCOVA test of the effects of the presence of epibionts and time-until-host-death on the infection intensity of three species of endoparasites.

| Endoparasite | Factor | F | p |
| --- | --- | --- | --- |
| *M. bicuspidata* | Presence of epibionts | 6.76 | **0.012** |
|  | Time-until-host-death | 85.12 | **<0.001** |
|  | Presence of epibionts × Time-until-host-death | 8.16 | **0.006** |
| *P. ramosa* | Presence of epibionts | 0.02 | 0.90 |
|  | Time-until-host-death | 25.44 | **<0.001** |
|  | Presence of epibionts × Time-until-host-death | 0.30 | 0.59 |
| *H. tvaerminnensis* | Presence of epibionts | 6.69 | **0.017** |
|  | Time-until-host-death | 0.02 | 0.89 |
|  | Presence of epibionts × Time-until-host-death | 1.14 | 0.30 |

Survival of *Daphnia* was mainly affected by the endoparasites. *Daphnia* exposed to *M. bicuspidata* had the lowest survival during the experiment, significantly lower than uninfected *Daphnia*, both with (Pairwise Wilcoxon: p<0.001) and without epibionts (Pairwise Wilcoxon: p<0.001). Survival of *Daphnia* infected by *P. ramosa* and *H. tvaerminnensis* was lower than the uninfected *Daphnia*, but the difference was not significant. In all groups the presence of epibionts did not affect the survival of the *Daphnia* (Figure 4).

**Figure 4:**
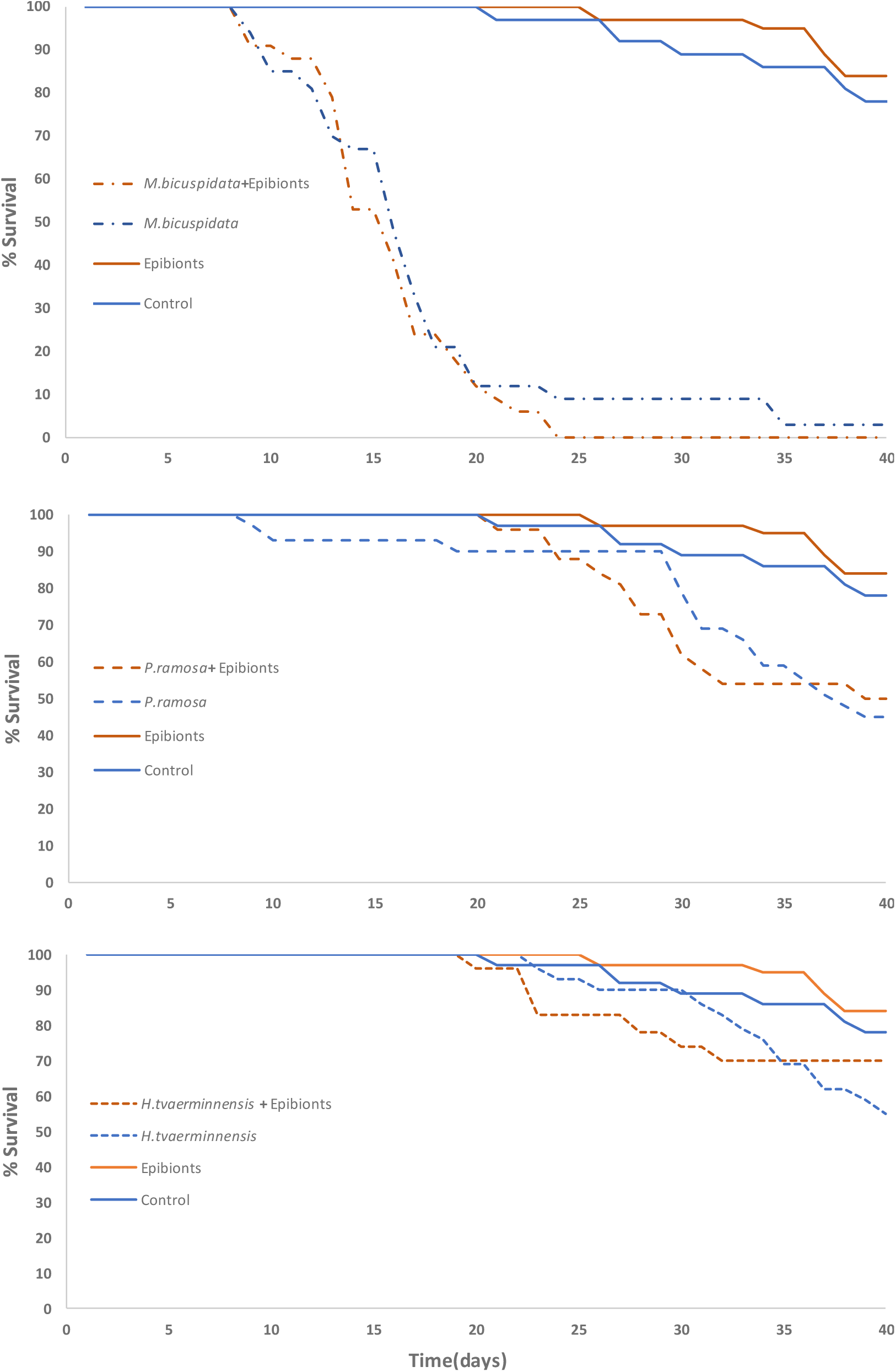
Survival of *D. magna* exposed to three species of endoparasites in the presence and absence of epibionts, including controls.

The number of offspring produced by the *Daphnia* host was affected by the presence and identity of the endoparasites (ANOVA F=37.19, p<0.001), but not by the presence of epibionts (ANOVA F=0.76, p=0.39) or the endoparasite-epibiont interaction (ANOVA F=1.4, p=0.27). More precisely, the number of offspring produced by the control group was significantly higher than the number of offspring produced by *Daphnia* exposed to *M. bicuspidata,* both with (Pairwise Wilcoxon: p<0.001) and without epibionts (Pairwise Wilcoxon: p<0.001). Similarly, the number of offspring produced by the control group was significantly higher than the number of offspring produced by *Daphnia* exposed to *P. ramosa*, both with (Pairwise Wilcoxon: p<0.001) and without epibionts (Pairwise Wilcoxon: p=0.006). However, the number of offspring produced by the control group was significantly higher than the number of offspring produced by *Daphnia* exposed to *H. tvaerminnensis* without epibionts (Pairwise Wilcoxon: p=0.031), but not with epibionts (Pairwise Wilcoxon: p=0.11). Furthermore, the number of offspring produced by *Daphnia* exposed to *H. tvaerminnensis* was significantly higher than the number of offspring produced by *Daphnia* exposed to *M. bicuspidata,* with (Pairwise Wilcoxon: p=0.002) and without epibionts (Pairwise Wilcoxon: p<0.001). The presence of epibionts did not affect offspring production in any of the groups (Figure 5).

**Figure 5:**
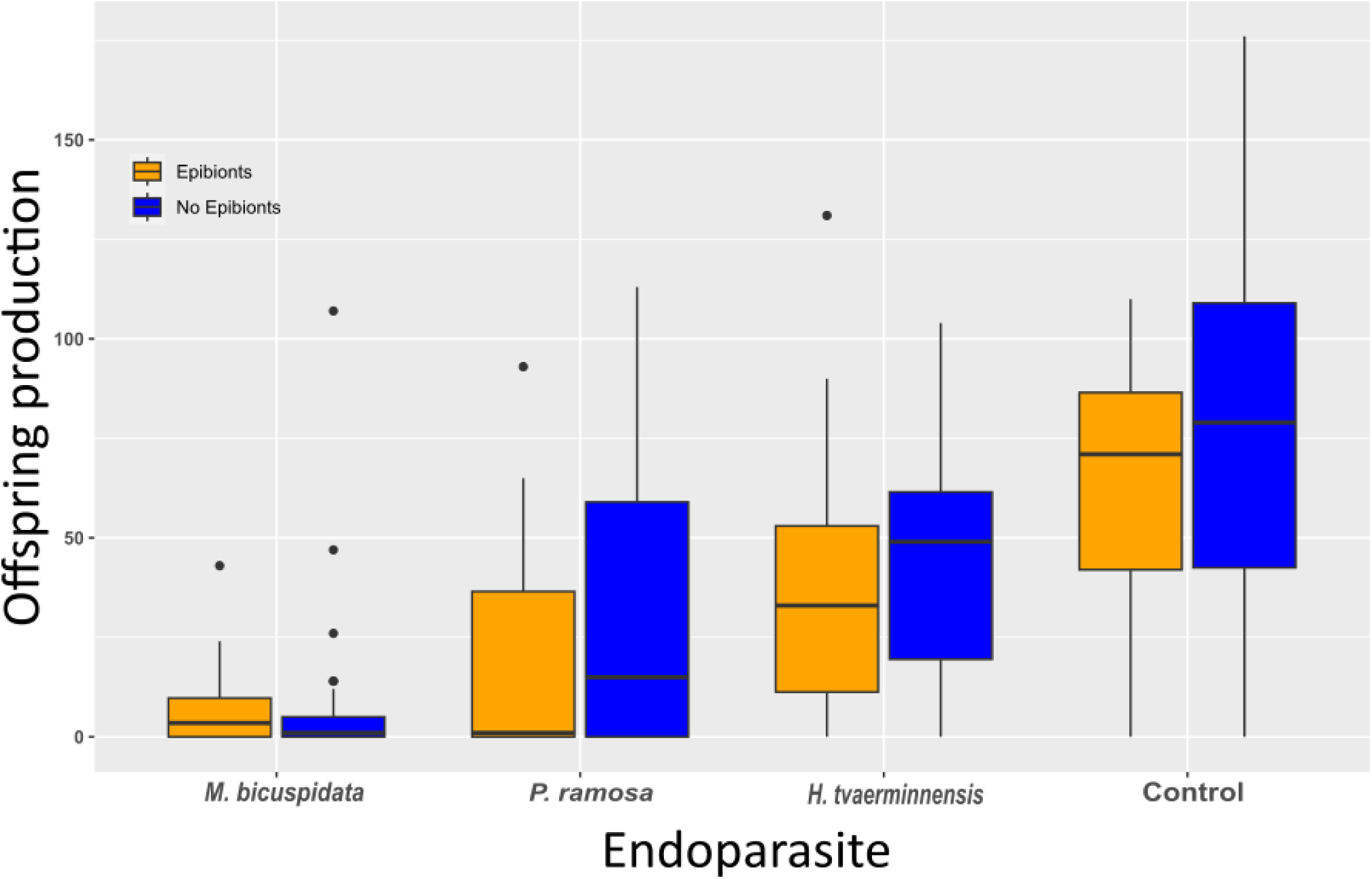
Mean±SE of the number of offspring produced by *D. magna* exposed to three species of endoparasites in the presence and absence of epibionts, including controls.

## Discussion

We conducted a series of experiments to study the relationship between endoparasites and epibionts and the effects of this relationship on their *Daphnia* host. We found that infection by two out of three species of endoparasites affected the attachment of epibionts, albeit in opposite directions, while infection by the third endoparasite did not affect epibiont attachment. We also found that the presence of epibionts affected the infection intensity of two out of three species of endoparasites, and this effect was not necessarily correlated with time-until-host-death. Host fitness was found to be affected by the presence and identity of endoparasites, but not by the presence of epibionts or by endoparasite-epibiont relationship.

The number of epibionts attached to *Daphnia* ranged between 0-17. In natural communities, *Daphnia* can become covered by dozens of epibionts (Iyer and Rao, 1993). Thus, the number of epibionts attached to *Daphnia* in our experiments seems rather low. These low numbers may be due to the number of epibionts placed in each jar in the beginning of the experiments (20 individuals) or short incubation time (6 hours). Our choice of these experimental settings is not without reason. First, when the number of epibionts is high, attachment space on the *Daphnia*’s body becomes scarce, resulting in competition over attachment space. Our aim was to exclude such competition, which may occur when *Daphnia* are covered by many epibionts. Second, when a *Daphnia* molts, it sheds the epibionts attached to it (Duneau and Ebert, 2012). Third, *Daphnia* may also produce offspring on which the epibiont may attach. Lastly, the *Daphnia* used in our experiments were already infected by one of the endoparasites, and thus they were likely to die during a long experiment. Short incubation time was thus required in order to avoid bias caused by molting, premature death or offspring production of the *Daphnia*. It should be noted that the difference in the number of attached epibionts between infected and uninfected *Daphnia*, despite being significant in most experiments, was relatively modest. This can be related to the relatively low number of attached epibionts.

Epibionts attached in different quantities to *Daphnia* infected by different species of endoparasites vs. uninfected *Daphnia*. It has been suggested that attachment of the epibiont *B. rubens* is affected by behavioral factors, such as the swimming velocity and frequency of antenna movements of *Daphnia* (Hirshberg and Ben-Ami, submitted). It is well known that the susceptibility to infection varies between young and adult hosts (reviewed in Ben-Ami, 2019), including in *Daphnia* (Izhar and Ben-Ami, 2015; Gattis et al., submitted), and that young and adult *Daphnia* exhibit different swimming behaviors (Gust et al., 2019). Moreover, different species of endoparasites may alter host behavior in different ways (Levri, 1999), such as depth selection and swimming activity. Thus, differences in behavior between infected and uninfected *Daphnia* could influence the attachment preference of the epibionts. Additionally, infection by endoparasites induces coloration changes of *Daphnia*, which may affect the predation preference of some of their predators (Goren and Ben-Ami, 2017; Johnson et al., 2006). The epibiont *B. rubens* is attracted towards its *Daphnia* host by the water currents it produces during swimming and filter feeding (May, 1989). Future work should focus on understanding the effects of endoparasites on the swimming and filter-feeding of *Daphnia* and their effect on epibiont attachment.

Epibionts attached more to uninfected *Daphnia* in comparison with *Daphnia* infected by *M. bicuspidata. M. bicuspidata* is highly virulent, killing its host within 2-3 weeks. Another fungal parasite, *Polycaryum leave*, was found to increase its host’s morbidity, reduce energy availability and thus decrease swimming movement (Johnson et al., 2018). Attaching to a moving *Daphnia* host was shown to be more beneficial to the epibiont *B. rubens* (Iyer and Rao, 1995), and in our cultures epibionts detached from the *Daphnia* when it was placed on a substrate on which it could not move. We selected *Daphnia* infected by *M. bicuspidata*, which may have decreased its host swimming behavior. This may explain the lower attachment of epibionts. In contrast to *M. bicuspidata,* epibionts attached more to *Daphnia* infected by *H. tvaerminnensis* than to uninfected *Daphnia*. Other microparasites were shown to affect depth selection of *Daphnia* (Fels et al., 2004). In our cultures, *B. rubens* was mainly seen at the bottom of the jars. Thus, here depth selection may have affected the attachment of epibionts between infected and uninfected *Daphnia*.

The epibiont rotifer competes with its *Daphnia* host over food resources (Pauwels et al., 2014). Given that decreasing the amount of food for the *Daphnia* host can decrease endoparasitic output, as the endoparasite depends on its host’s resources for growth and reproduction (Manzi et al., 2020), we expected that the presence of the epibiont would decrease the amount of food available for the *Daphnia* and its endoparasites, reducing infection prevalence and intensity. Nevertheless, the prevalence and intensity of infection in our experiments was found to be equal or higher in the presence of epibionts. Because our experimental *Daphnia* were fed daily with high-quality food, we conjecture that the amount of food lost to the epibionts may have been negligible, still enabling the growth and reproduction of both *Daphnia* and its endoparasites.

Infection prevalence of endoparasites often correlates with the concentration of endoparasitic transmission stages, i.e., dose effect (Ben-Ami et al., 2008; Pulkkinen, 2007). *Daphnia* have been found to reduce their filter-feeding rate in the presence of high density of conspecifics (Pulkkinen, 2007). Thus, the presence of competitors may negatively affect the filtering rate of *Daphnia*. On the other hand, *Daphnia* were found to increase their filtration rate when covered by epibiont algae (Allen et al., 1993). Being both a competitor and an epibiont, *B. rubens* may affect the *Daphnia*’s filter-feeding in several ways. Changes in the *Daphnia*’s filtration rate may affect the number of endoparasite spores entering the *Daphnia* abdomen and thus affect infection. Rotifers of the genus *Brachionus* are filter-feeders, filtering small particles and cells from the surrounding water (Gilbert, 2022). In our experiments, spores of *P. ramosa* were found in the gut of *B. rubens.* If the epibiont *B. rubens* indeed consumes endoparasitic spores, it would have decreased infection prevalence rather than increasing it (i.e., dilution effect), as was seen in our experiments. Possibly, the endoparasitic spores remain viable and capable of infecting the *Daphnia* after passing through the epibiont’s digestive system, as was found in the case of *P. ramosa* (King et al., 2012). Filter feeding by the epibionts may have also created water currents, causing the endoparasitic spores to flow at the water column rather than sink to the bottom of the jar, thereby increasing encounter between endoparasitic spores and *Daphnia*, and resulting in higher infection prevalence.

When attached to their host’s carapace, epibionts may harm its buoyancy and mobility (Pauwels et al., 2014). On the one hand, harming host buoyancy may cause it to sink to the bottom of the jar (or to the sediment in natural habitats; Barea-Arco et al., 2001), exposing it to higher concentrations of endoparasitic spores, which may increase infection prevalence. On the other hand, disruption by epibiont burden may cause the host to spend more resources on swimming, reducing the resources available for endoparasite reproduction and reducing its infection intensity. The high quantity of food given to the *Daphnia* in our experiments may have provided sufficient energy for both *Daphnia* swimming and endoparasite reproduction.

Previous studies of the *Daphnia*-microparasites system have revealed strong genotype-genotype interactions (Carius et al., 2001; Orlansky & Ben-Ami, 2019). Thus, different *Daphnia* genotypes vary in their susceptibility to different endoparasite genotypes (Luijckx et al., 2011). The *Daphnia* clone used in our experiments was shown to be susceptible to all endoparasite species as well as to the epibiont. Repeating the experiments using different genotypes of the *Daphnia,* endoparasites and epibionts may produce different insights.

In a series of competition experiments, Gilbert (1985) found that rotifer populations decline in the presence of *Daphnia*, mainly as a result of exploitative competition, which affects the rotifers but not the *Daphnia*. In our experiments, new rotifers were added to the jars after 10 and 30 days, due to their population decline. Competitive exclusion of the epibiont may explain their lack of influence on *Daphnia* survival and reproduction. Both survival and reproduction of *Daphnia* were negatively affected by the endoparasites. Of the three species of endoparasites, *M. bicuspidata* was the most infectious and virulent, infecting almost 100% of the *Daphnia* and killing them within 9-25 days. The low offspring production of *Daphnia* exposed to *M. bicuspidata* is likely due to their fast death. The bacterium *P. ramosa* castrates its host before killing it within 30-50 days (Ebert et al., 1998), which (for a different reason) explains the low offspring production of *Daphnia* exposed to *P. ramosa*. *H. tvaerminnensis* is transmitted both vertically (from mother to offspring) and horizontally (via water born environmental spores) (Haag et al., 2011; Orlansky and Ben-Ami, 2019; Orlansky and Ben-Ami, 2023). Thus, the transmission of *H. tvaerminnensis* depends on the reproduction success of its host, which explains its relatively negligible influence on *Daphnia* survival and reproduction.

Infection intensity (parasite spores produced) of *M. bicuspidata* and *P. ramosa* correlated with time-until-host-death. In other words, the longer the host lived, the more time endoparasites had to reproduce within its body. Additionally, the longer *Daphnia* lived, the larger it became (including the manifestation of gigantism), and larger *Daphnia* can support larger population of endoparasites (Ebert et al., 2004). This apparent benefit for the parasite does not come without a cost if, for example, the host is predated (Ben-Ami, 2017; Acevedo et al., 2019). Interestingly, the infection intensity of *H. tvaerminnensis* did not correlate with time-until-host-death, with most *Daphnia* exposed to *H. tvaerminnensis* surviving for 40 days until the end of the experiment. Although prolonging the experiment might have revealed that the infection intensity of *H. tvaerminnensis* also correlates with time-until-host-death, it is likely that mixed-mode transmission also influenced the virulence-transmission trade-off (Ebert, 2013).

Epibiont-endoparasite relationships have been studied in various host taxa. For example, when studying willow ptarmigan, Holmstad et al. (2008) found a correlation between epibiont load and endoparasitic infection. They hypothesized than endoparasitic infection reduces the host energy available for peering against the epibionts. Balestrieri et al. (2006) found that the ectoparasite *Sarcoptic mange* and endoparasitic helminths co-occurred on wild foxes. Additionally, James et al. (2002) found evidence showing that sheep with greater resistance to gastrointestinal parasites are less susceptible to lice. In contrast, when studying the trout *Salmo trutta,* Larsen et al. (2002) found a negative relationship between the endoparasite *Anisakis* sp. and the ectoparasite *Gyrodactylus derjavini,* possibly due to the activation of the fish’s immune system. These results emphasize the need for studying the interactions between endoparasites and epibionts, interactions whose results are crucial for understanding the dynamics of planktonic communities.

## Conclusions

Endoparasites and epibionts constitute important components of natural zooplankton communities, as they are widely spread and infect a variety of hosts. Epibionts were found to attach in different quantities to hosts infected by different species of endoparasites. By attaching to a cladoceran host, the epibiont benefits in several ways, including defense from predation, energy saving and access to new food sources. By changing the host’s availability to the epibiont, the endoparasite may affect its population abundance and survival in the waterbody. Despite their small effect on host survival and fecundity, epibionts were found to increase infection prevalence and intensity of several endoparasites, most likely due to the endoparasite’s virulence and transmission mode. Thus, the attachment of epibionts may increase the prosperity of endoparasites in aquatic communities, and by increasing endoparasitic infection, epibionts may indirectly (and inadvertently) change the composition of cladoceran communities.

## Acknowledgements

We are grateful to E. Marcus, T. Hiskia and N. Barbarshteyn for their assistance in the preforming of the experiments.

## Data availability

Data supporting this study is available on figshare, DOI 10.6084/m9.figshare.24922614.

## Author contribution

OH and FBA conceived and designed the study. OH performed the research, analyzed the data and wrote the paper. FBA obtained funding and supervised this study, and provided editorial advice. All authors read and approved the final version of the manuscript.

